# TXNRD3 supports male fertility via the redox control of spermatogenesis

**DOI:** 10.1101/2021.12.29.474493

**Authors:** Qianhui Dou, Anton A. Turanov, Marco Mariotti, Jae Yeon Hwang, Huafeng Wang, Sang-Goo Lee, Joao A. Paulo, Sun Hee Yim, Stephen P. Gygi, Jean-Ju Chung, Vadim N. Gladyshev

## Abstract

Thioredoxin/glutathione reductase (TGR, TXNRD3) is a thiol oxidoreductase of unknown function composed of thioredoxin reductase and glutaredoxin domains. This NADPH-dependent enzyme evolved by gene duplication within the *Txnrd* family, is expressed in the testes and can reduce both thioredoxin and glutathione *in vitro*. To characterize the function of TXNRD3 *in vivo*, we generated a strain of mice with the deletion of *Txnrd3* gene. We show that *Txnrd3* knockout mice are viable and without discernable gross phenotypes, but TXNRD3 deficiency leads to fertility impairment in male mice. *Txnrd3* knockout animals exhibit a lower fertilization rate *in vitro*, a sperm movement phenotype and an altered redox status of thiols. Proteomic analyses revealed a broad range of substrates reduced by TXNRD3 during sperm maturation, presumably as a part of quality control. The results show that TXNRD3 plays a critical role in male reproduction via the thiol redox control of spermatogenesis.

## Introduction

Thioredoxin reductases (TXNRDs) are members of the pyridine nucleotide disulfide oxidoreductase family, which utilize NADPH for the reduction of their thioredoxin substrates. Three TXNRD paralogs are present in mammals: TXNRD1, TXRND2, and TXRND3 (1). These proteins have a C-terminal penultimate selenocysteine (Sec) residue inserted co-translationally in response to UGA codon. TXNRD1 and TXRND2 are thioredoxin reductases in the cytosol and mitochondria, respectively, and are ubiquitously expressed in various tissues and cells (2). Previous studies revealed that TXNRD1 and TXNRD2, like their substrates (cytosolic Trx1 and mitochondrial Trx2), are essential enzymes, i.e. their knockout leads to early embryonic lethality in mice (3, 4). These and other studies revealed a critical role of TXNRDs and more generally of thiol-based redox control in mammals.

The third mammalian thioredoxin reductase, TXNRD3, is unusual in that it has an additional N-terminal glutaredoxin (Grx) domain, which is fused to the canonical TXNRD module. Due to this unusual domain organization, this enzyme can reduce both thioredoxin and glutathione (GSH), and therefore was termed thioredoxin/glutathione reductase (5). GSH and Trx pathways are two major pathways that maintain the reduced state of cellular thiols and regulate redox homeostasis (6), and the fact that one enzyme can support the function of both pathways is of great interest. Like other TXRNDs, TXNRD3 is a homodimer, with the monomers oriented in a head-to-tail manner. In mammals, TXNRD3 expression is largely restricted to the testis, and our previous studies suggested that TXNRD3 may be involved in the process of sperm maturation (7). It was proposed that TXNRD3 supports disulfide bond formation and isomerization. Interestingly, in platyhelminth parasites, TXNRD3 is the only TXNRD encoded in the genome, and it was found that this enzyme maintains the reduced state of GSH and Trx in both cytosol and mitochondria (6).

In the current work, to dissect the role of TXNRD3 in redox homeostasis and male reproduction *in vivo*, we prepared mice deficient in *Txnrd3*. In contrast to *Txnrd1* and *Txnrd2* knockout mice, *Txnrd3* knockout mice are viable, but the deletion of this gene leads to male fertility impairment. We show that TXNRD3 functions to support the reduction of its substrates during epididymal sperm maturation. Accordingly, TXNRD3 deficiency disrupts redox homeostasis during this process, leading to abnormal motility of the sperm and ultimately to suboptimal male fertility.

## Materials and Methods

### Gene targeting, mouse breeding and genotyping

*Txnrd3* knockout (KO) mice were generated using a gene targeting strategy that replaced the third coding exon of *Txnrd3* with an IRES/bGeo/PolyA cassette via homologous recombination. Mutant ES clones were generated and confirmed by Southern blotting with 5’ and 3’ flanking probes. The Southern probes were as follows: 5’ external probe 1, 5’-CTCCTCAAGGCAGTCATAACAC, and 5’ external probe 2, 5’-AGCACACCCTGGCCAACTTACC, generated a 7.6 kb product of the wild type allele and a 174 kb product of the mutant allele; 3’ internal probe 1, 5’-CCTCAGAAGAACTCGTCAAG, and 3’ internal probe 2, GGCAGCGCGGCTATCGTG, produced a 9.8 kb product in the mutant allele only. Chimeric mice were then generated by the injection of ES cells into blastocysts of C57BL/6 mice. *Txnrd3*^-/-^ mice were obtained by crossing heterozygous progeny of chimeric mice. Genotyping was performed using genomic DNA isolated from mouse tail biopsies using allele-specific primers as follows: primer 19, 5’-CAGGTGTCCTCTGAAGGCATC, primer 20, 5’-CACAGTGTTGAGGACCGTGTC, were designed for the detection of the wild type allele, generating a 322 bp band; primer 19, and primer GT-IRES, 5’-CCCTAGGAATGCTCGTCAAGA, were designed for the detection of the mutant allele, producing a 441 bp band.

Mice were kept under standard conditions with food and water. All animal experiments were performed in compliance with the recommendations in the Guide for the Care and Use of Laboratory Animals of the National Institutes of Health and have been approved by the Institutional Animal Care and Use Committees (IACUCs) of Brigham and Women’s Hospital and Yale University.

### Preparation of mouse spermatozoa

Adult male mice (8-10 weeks old) were asphyxiated, and the epididymis was removed. Spermatozoa were isolated from the caput, corpus, and caudal regions of the epididymis. Caudal epididymal mouse sperm was collected by swim-out in M2 media, allowing motile sperm to disperse for 15 min at 37 °C and removing connective tissues. Suspended cells were harvested into a microtube, and the non-sperm cells and immotile sperm cells were let sediment for 5 min. Sperm cells were diluted 20-fold in media and counted with a hemocytometer.

### Sperm motility analysis

Caudal epididymal spermatozoa were suspended and incubated in a non-capacitating M2 medium (Millipore). Uncapacitated spermatozoa were allowed to disperse for 15 min at 37 °C in M2 media, and free-swimming sperm cells were recorded with a microscope during the next 5 min.

### Fertility tests and *in vitro* fertilization

To cross heterozygous mice, two females were caged with one male, and their pregnancy and litter production were tracked. For a mating experiment across genotypes (WT x WT, heterozygous KO x heterozygous KO, and homozygous KO x homozygous KO), one female was caged with one male, with 7 pairs of mice in each group. After 2 weeks, female mice were separated from males and their pregnancy and litter production were tracked. In vitro fertilization (IVF) was performed as previously described with minor modifications (8). Four-six-weeks old B6D2F1 female mice were subjected to superovulation by serial injection of progesterone, anti-inhibin serum (Central Research Co, Ltd), and human chorionic gonadotrophin (hCG, EMD Millipore) (9). The oocytes were collected 13 hours after hCG injection. Prepared cauda sperm cells were capacitated *in vitro* at 37 °C, 5% CO_2_ for 1 hour in human tubular fluid (HTF, EMD Millipore) at 2.0 × 10^6^ cells/ml and co-incubated with oocytes at 2.0 × 10^5^ sperm/ml. After 4.5 h of the co-incubation, the oocytes were washed and transferred to a fresh HTF medium. Following overnight culture at 37 °C under 5% CO_2_, 2-cell embryos were counted to measure the fertilization rate.

### *Txnrd3* gene expression analyses

Gene expression data was collected from the EBI gene expression atlas (https://www.ebi.ac.uk/gxa/home), selecting datasets of tetrapods for which testis data was available. All samples were from adult males. The analyzed species included human (GTEx (10)), mouse (Fantom5 (11)), rat (12), sheep (13), cow, chicken, and macaque (14).

### Immunohistochemistry

Testes and epididymis were fixed in 10% neutral buffer formalin overnight and dehydrated with graded concentrations of alcohol prior to being embedded in paraffin. Paraffin-embedded tissues were sectioned into 4 *μ*m slices and placed on slides. Immunohistochemistry staining was performed on paraffin-embedded tissue sections. We used the following in-house primary antibodies: TXNRD3 (1:200), GPX4 (1:200), and PRDX4 (1:200). Staining was visualized using Goat anti-Rabbit IgG conjugated with HPR polymer and 3,3-diaminobenzidine (DAB) (Vector lab). Slides were scanned on an Axio Scan.Z1 (Zeiss) and the whole mount digitalized at 10x magnification.

### Immunoprecipitation and western blotting

For immunoprecipitation, 1 × 10^7^ sperm cells were sonicated briefly in 1% Triton X-100 in PBS with a protease inhibitor cocktail, further extracted for 4 h at 4 °C and cleared by centrifugation at 10,000 x g for 30 min. Testis was lyzed in a cell lysis buffer for 10 min. Solubilized proteins in the lysates were immunoprecipitated with an anti-GSH antibody and stained with the indicated antibodies.

For Western blotting, we added ice-cold RIPA lysis buffer to mouse testis tissue, homogenized using an electric homogenizer, and agitated the material for 2 h at 4°C. Mouse epididymal spermatozoa were washed in PBS, and 1×10^6^ sperm cells were directly lysed in a 2×SDS sample buffer. The whole sperm lysate and the testis lysate were centrifuged at 15,000 g, 4°C for 10 min. Supernatant was treated with 50 mM DTT and denatured at 95°C for 10 min prior to loading onto a gel. Antibodies used for Western blotting were TNXRD3 (1:1000), GPX4 (1:500), PRDX4 (1:500), GSH (1:1000), TXN1 (1:500), GSTA2 (1:500), GSTM1 (1:1000), and β-actin (1:1000). Secondary antibodies were anti-rabbit IgG-HRP (1:10,000) and anti-mouse IgG-HRP (1:10,000).

### Flow cytometry

For ROS measurements, caudal epididymal mouse sperm was incubated in PBS containing 5 mM CM-H2DCFDA at 37 °C for 30 min in the dark, then suspended in PBS buffer for flow cytometry analyses. To assay thiols, caudal epididymal mouse sperm was incubated in PBS containing 50 μM of Bodipy-NEM for 30 min in the dark, then suspended in PBS for flow cytometry analysis. 10,000 cells were analyzed with a FACSCalibur flow cytometer (Becton Dickinson). Data was analyzed using FlowJo software.

### Diagonal two-dimensional analyses

Electrophoresis on 2D gels was performed as previously described (15). Four independent runs were conducted of the samples from *Txnrd3* KO testis lysate, WT testis lysate, *Txnrd3* KO sperm lysate, and WT sperm lysate. Gel images were acquired with an Image Scanner III.

### Proteomic analyses

Aiming to identify TXNRD3 target proteins, we collected luminal content of caput, corpus, and cauda regions of the epididymis of *Txnrd3* KO and WT mice, pulling material from 8 mice to yield the amounts sufficient for further analyses. We subjected the samples to a procedure designed to enrich endogenous target proteins (Fig. S4). Our strategy assumed that target proteins are in an oxidized form in the *Txnrd3* KO testis. We prepared a thiol-trapping column by adding 5 ml homogenous suspension of resin onto the column equilibrated with the flow buffer, washed the resin with 5-10 volumes of H2O, and then with 5-10 column volumes of a binding buffer (50 mM Tris-HCl, pH 8.0, 100 mM NaCl, 0.1 mM EDTA). The luminal content including sperm from each epididymal region was lysed with RIPA buffer, supplemented with 50 mM N-ethylmaleimide (NEM) for 5 min, and applied onto PD MidiTrap G-25 column to clean up the excess of NEM prior to loading the samples. Under these conditions, NEM permanently modified reduced cysteines in proteins. In *Txnrd3* KO mice, TXNRD3 targets would still be partially oxidized, and these oxidized residues would not react with NEM. In order to specifically reduce TXNRD3 target proteins, we generated a column with an immobilized reduced Grx-domain of TXNRD3. We loaded the purified Grx domain onto a His Trap column, added 5 ml binding buffer containing 10 mM fresh DTT to reduce the domain, and then applied the epididymal luminal lysate and collected the eluate. Finally, the eluate was immediately applied to a pre-prepared thiol-trapping column to trap specifically reduced cysteines, and the bound proteins were eluted with 5 column volumes of the elution buffer (50 mM Tris-HCl, pH 8.0, 300 mM imidazole, 50 mM NaCl, 0.1 mM EDTA) containing 5 mM DTT. The eluate from this step was alkylated with 50 mM iodoacetamide (IAM) and subjected to LC-MS/MS for protein identification.

### Statistical analyses

Analysis was performed using GraphPad Prism 8. Data analyses were made using unpaired t tests. Statistical significance was denoted by p < 0.05. Errors are reported as standard error of means (SEM) unless otherwise indicated.

## Results

### Evolution of the TXNRD family to a conserved testis-specific TXNRD3

In mammals, TXNRDs are selenoproteins containing a C-terminal penultimate selenocysteine (Sec), the 21^st^ amino acid encoded by UGA. Compared with the 55 kDa subunit homodimeric TXNRD1 or TXNRD2, TXNRD3 is composed of two 65 kDa subunits because it has an additional N-terminal glutaredoxin (Grx) domain (Fig. 1*A*). The reductase function of TXNRDs is supported by a conserved CxxxC motif, which accepts electrons from FAD thereby mediating the flow of reducing equivalents from NADPH to Trx (Fig 1*B*). Our analysis (7) suggested that mammalian TXNRD1 and TXNRD3 evolved by gene duplication from an ancestral protein similar to fish Grx-containing TXNRD1 (Fig. 1*C*). TXNRD1 then lost the Grx domain (except for some rare isoforms), whereas the second Cys in the catalytic CxxC motif of the Grx domain of TNXRD3 was mutated to serine. We examined the average expression of TNXRD3 mRNA (transcripts per million (TPM)) across different tissues of various species of tetrapods. Unlike TXNRD1 (present in the cytoplasm in various tissues) and TXNRD2 (present in mitochondria in various tissues), in most analyzed species TXNRD3 is highly expressed only in the testis (Fig. 1*D*). This observation is strongly suggestive of a role of TXNRD3 in male reproduction, as well as that this biological function is conserved across tetrapods.

**Fig. 1.**
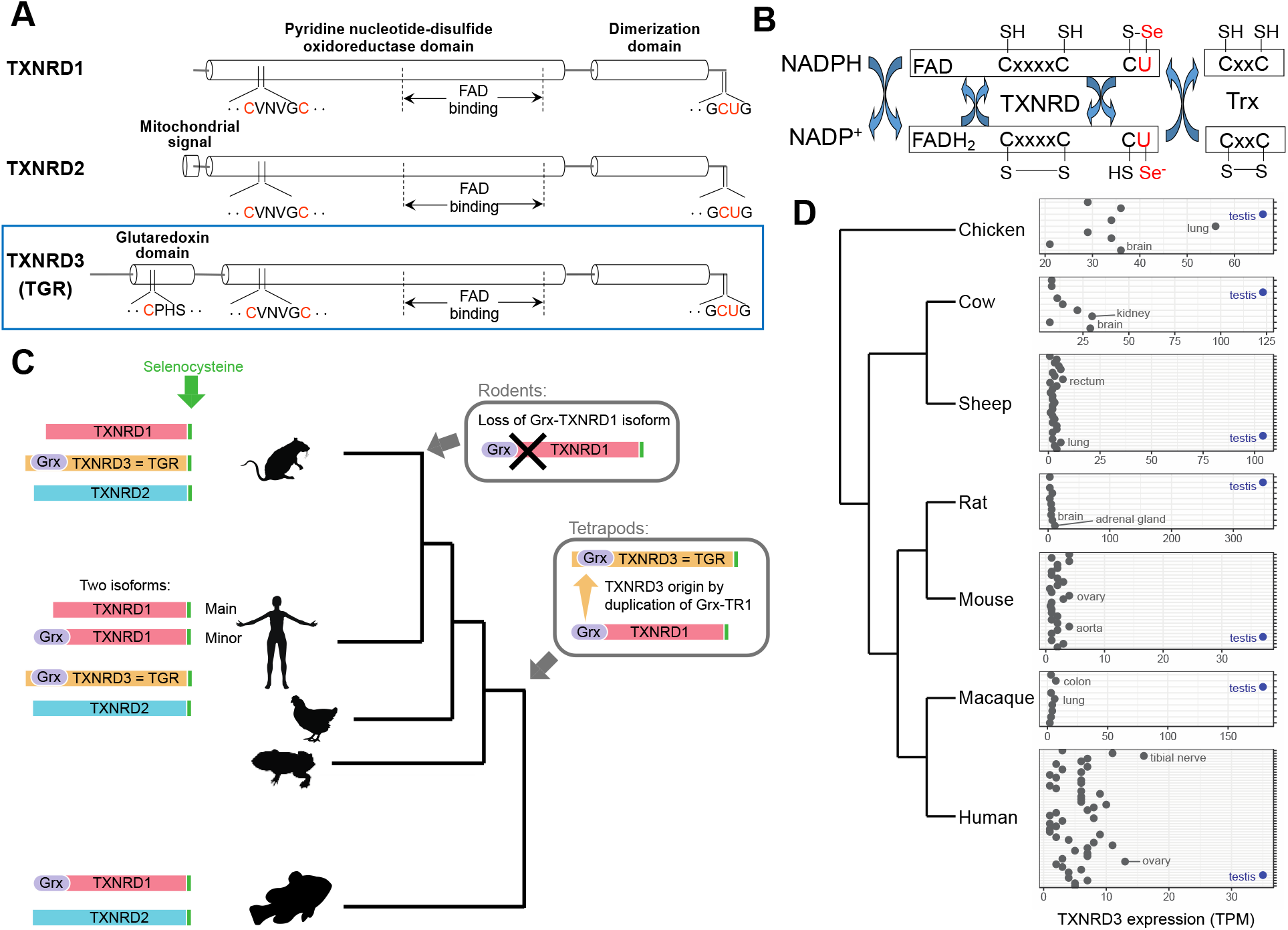
Organization and evolution of TXNRD3. (A) Structure of thioredoxin reductase (TXNRD) family members. (B) Flow of reducing equivalents in TXNRDs. The Grx domain of TXNRD3 may take place of Trx in accepting reducing equivalent from selenocysteine residue (U). (C) Evolution of TXNRD family in tetrapods. (D) Expression of TXNRD3 across tissues of tetrapods. Different sets of tissues are displayed per species, depending on data availability. For each species, the three tissues with highest TXNRD3 expression levels are labelled. Testis, which is the tissue with top expression for all species, is colored in blue.

### TXNRD3 deficient mice are viable but exhibit impaired fertility

To get insights into the function of TXNRD3 *in vivo*, we generated *Txnrd3* KO mice. The murine *Txnrd3* gene comprises 7 exons, and the *Txnrd3*-deficient mice were generated using a targeting strategy that replaced the third exon of *Txnrd3* (Fig. S1*A*). ES cell clones were tested for homologous recombination by PCR and verified by Southern blotting (Fig. S1*B*). Upon injection into C57BL/6 blastocysts, a recombinant ES clone contributed to chimeric males which transmitted the targeted allele to the germ line. The resulting *Txnrd3*-deficient mice were genotyped using mutant and allele-specific primers (Fig. S1*C*).

This procedure yielded viable homozygous *Txnrd3* KO mice, indicating that TXNRD3, in contrast to two other mammalian TXNRDs, is not essential during embryonic development and subsequent life. We also observed no other gross phenotypes in *Txnrd3* KO mice, in both males and females. Since the *Txnrd3* knockout allele was present in germ cells, its F1 x F1 cross should result in offspring having all three possible genotypes (*Txnrd3* ^+/+^; *Txnrd3* ^+/-^; *Txnrd3* ^-/-^) with the Mendelian ratio of 1:2:1. However, the actual ratio was 1:1.7:0.56 (WT: heterozygous: homozygous) (Table S1). This reduction in the number of homozygous mutants suggested a possibility of fertility impairment involving TXNRD3-deficient germline. To quantify this phenotype, we set up three mating groups of mice, including WT x WT, heterozygous KO x heterozygous KO, and homozygous KO x homozygous KO, with 7 pairs of animals in each group. By keeping males with females for 2 weeks and later recording the timing of females giving birth, we found that it took longer for *Txnrd3*-null females to give birth to offspring (WT 24.43±4.54 days vs. *Txnrd3*^-/-^ 36.57±16.27 days, p=0.081, n=7; WT 24.43±4.54 days vs. *Txnrd3*^+/-^ 35.00±17.18 days, p=0.141, n=7; *Txnrd3*^+/-^ 35.00±17.18 days vs. *Txnrd3*^-/-^ 36.57±16.27 days, p=0.863, n=7). It is also of note that two *Txnrd3*^-/-^ females did not give birth (they monitored for 60 days) (Fig. 2*A*), supporting a potential role of TXNRD3 for the germline. Additionally, there were fewer pups per litter in the case of homozygous KO mice (WT 7.43±1.72 vs. *Txnrd3*^-/-^ 3.71±2.69, p=0.010, n=7; WT 7.43±1.72 vs. *Txnrd3*^+/-^ 4.29±2.98, p=0.033, n=7; *Txnrd3*^+/-^ 4.29±2.98 vs. *Txnrd3*^-/-^ 3.71±2.69, p=0.713, n=7) (Fig. 2*B*).

**Fig. 2.**
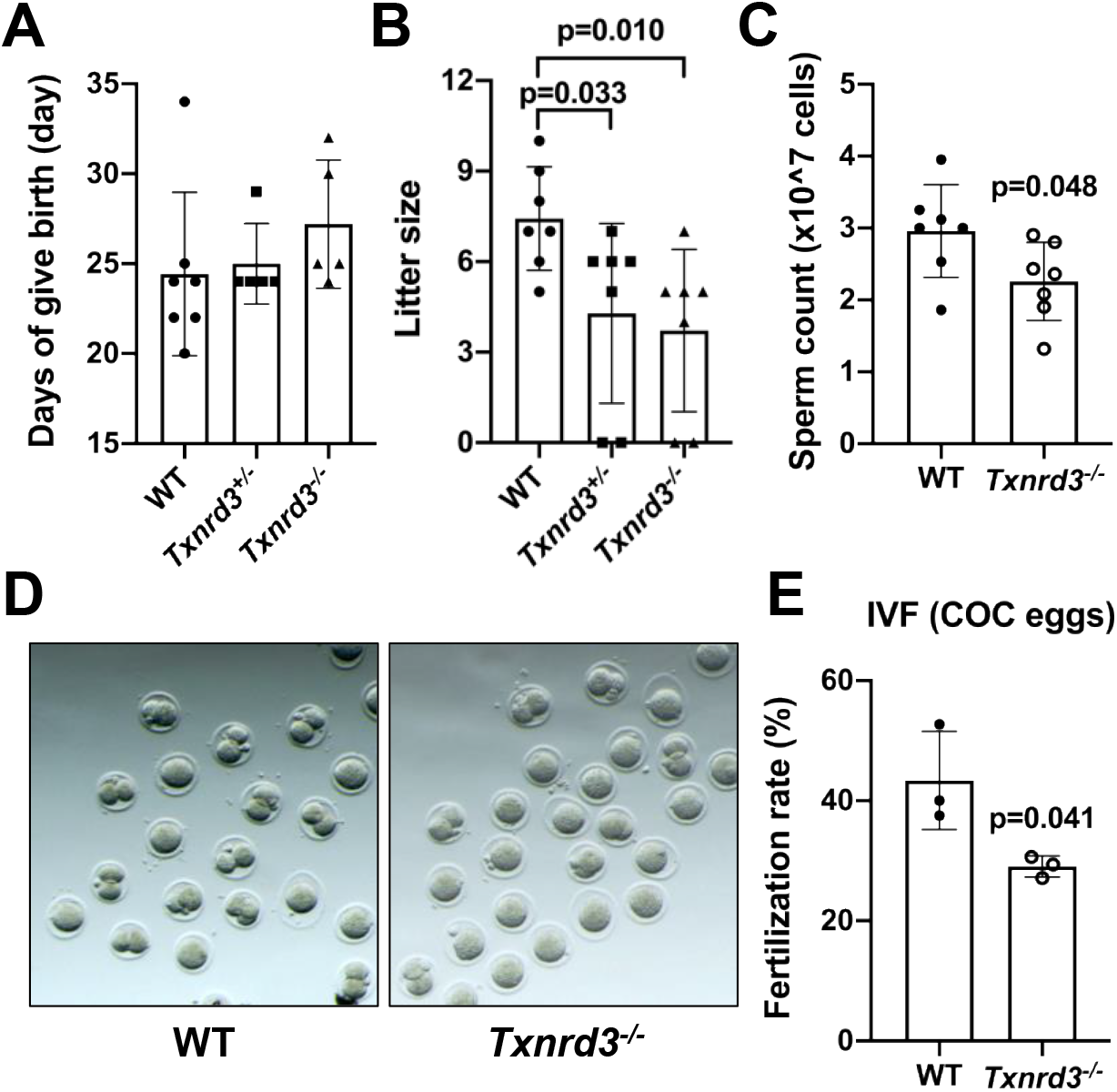
Deletion of mouse *Txnrd3* gene impairs male fertility. (A) Period from the beginning of mating to females giving birth. Two females in the heterozygous and homozygous groups did not give birth for 60 days after the beginning of mating. Hetero, heterozygous group, WT, wild type group, KO, homozygous group. (B) Litter size in each animal group. (C) Number of cauda sperm in WT and *Txnrd3*-null male mice. (D) Representative image of fertilized embryos from the cauda sperm from WT (*left*) and *Txnrd3*-null (*right*) males *in vitro*. (E) Fertilization rates of WT and *Txnrd3*-null animals *in vitro*. *, p>0.05.

We further examined sperm counts of *Txnrd3*-null and WT mice and found that the number of spermatozoa in *Txnrd3*-null mice was significantly lower than in WT mice (WT 2.96±0.64 × 10^7^ vs. *Txnrd3*^-/-^ 2.26±0.55 × 10^7^, p=0.048, n=7) (Fig. 2*C*). To further explore fertility of *Txnrd3*-null mice, we carried *in vitro* fertilization. The *in vitro* fertilization rate with COC-intact eggs was significantly lower when the sperm from *Txnrd3*^-/-^ males was compared to the WT sperm (WT 43.40±8.15 vs. TGR^-/-^ 29.03±1.78, p=0.041, n=3) (Fig. 2, *D* and *E*). Finally, we examined movement of free-swimming spermatozoa isolated from WT, *Txnrd3*^+/-^, and *Txnrd3*^-/-^ mice. Compared to WT mice, the fraction of motile spermatozoa was visibly lower in *Txnrd3*^-/-^ mice (Movie S1).

Taken together, our data suggest that while TXNRD3 is neither essential for development nor for male fertility in general, its KO leads to reduced sperm motility and fertility, making these male subfertile. In the accompanying manuscript (16), we provide further evidence for the role of TXNRD3 in epididymal maturation, and specifically its impact on capacitation-associated mitochondrial activation and sperm motility.

### Thiol peroxidases as interacting partners of TXNRD3

Previous functional characterization and localization of TXNRD3 revealed its possible disulfide bond isomerization activity and a close functional relationship to another selenoprotein, GPX4 (7). The use of String to examine the association between TXNRD3 and GPX4 showed a combined score of 0.717, suggesting a high confidence functional link between these two proteins (Fig. S2*A*). To further characterize this association, we analyzed protein levels of TXNRD3 and GPX4 in the mouse testis and sperm by immunoblotting. Interestingly, GPX4 expression was higher in *Txnrd3*^-/-^ testes than in WT testes (WT 0.77±0.11 vs. *Txnrd3*^-/-^ 0.96±0.04, p=0.049, n=3; WT 0.77±0.11 vs. *Txnrd3*^+/-^ 0.82±0.02, p=0.424, n=3; *Txnrd3*^+/-^ 0.82±0.02 vs. *Txnrd3*^-/-^ 0.96±0.04, p=0.006, n=3) (Fig. 3*A*), whereas the expression of this protein in the sperm was not much affected (Fig. 3*B*). Immunohistochemical staining of TXNRD3 and GPX4 showed a similar pattern (Fig. 3, *C* and *D*).

**Fig. 3.**
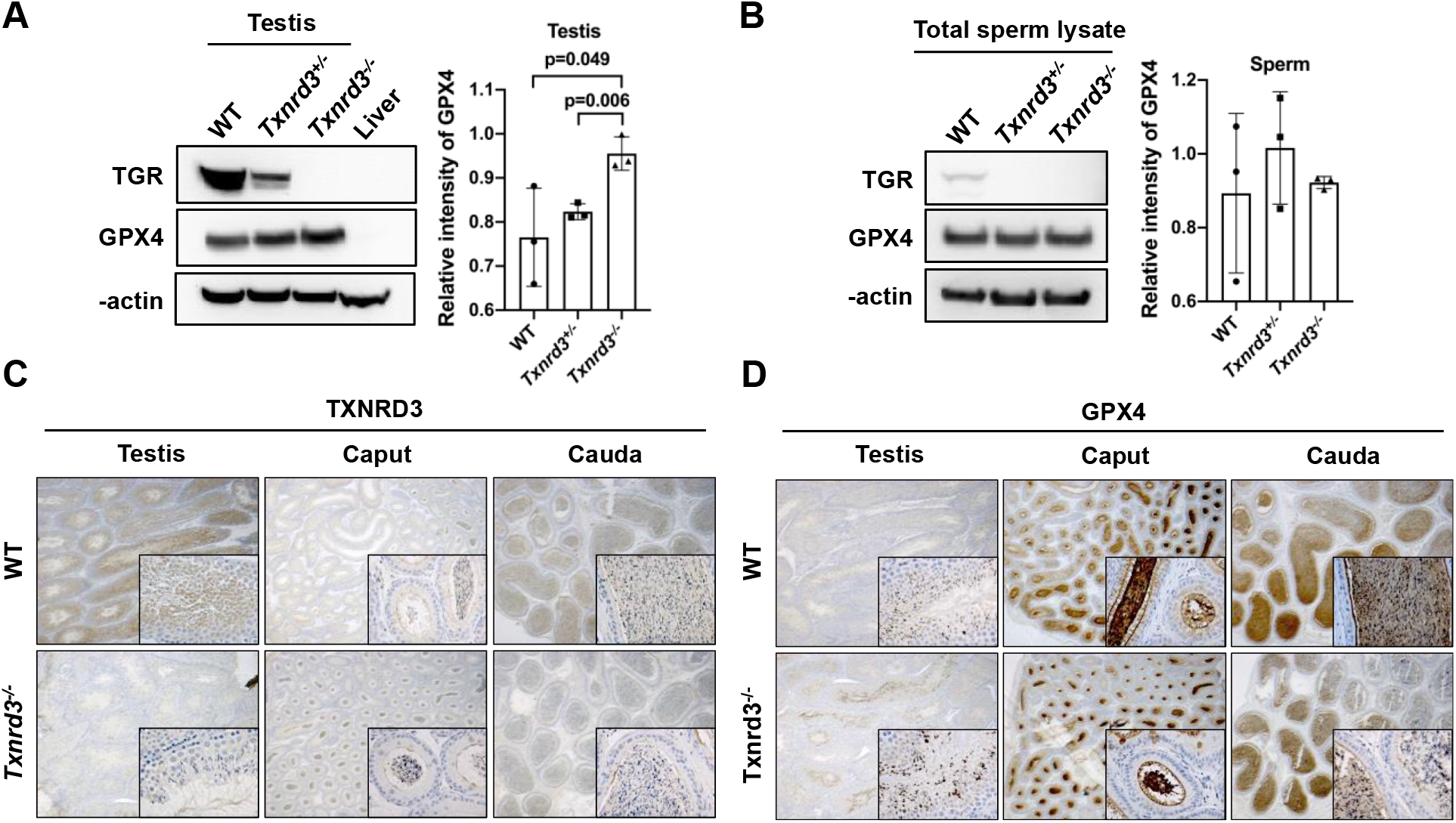
Expression of TXNRD3 and GPX4 in mouse testis and sperm. (A) TXNRD3 and GPX4 expression in testis based on Western blotting. The right panel quantifies GPX4. (B) TXNRD3 and GPX4 levels in mouse sperm. The right panel quantifies GPX4. (C) Immunohistochemistry of GPX4 in *Txnrd3* KO and WT mouse testis and sperm (caput and cauda). (D) Immunohistochemistry of TXNRD3 in *Txnrd3* KO and WT mouse testes and sperm (caput and cauda).

Another group of thiol peroxidases implicated in male reproduction are peroxiredoxins (PRDXs) (17). In particular, peroxiredoxin 4 (PRDX4) is thought to play a role in this process by regulating oxidative packaging of sperm chromatin (18). Based on String, the association among PRDX4, GPX4, and TXNRD3 showed a medium confidence (Fig. S2*A*). We detected the expression of PRDX4 in mouse testis and sperm, and its expression in the testis was particularly high. We also found that PRDX4 expression was decreased in *Txnrd3*-deficient testis compared to WT (Fig. S2, *B* and *C*). Altogether, these analyses were suggestive of a possible functional association of TXNRD3 with thiol peroxidases of the GPX and PRDX families, but further studies are needed to clarify this possibility.

### A role of Txnrd3 in maintaining thiol redox status of sperm proteins

Redox homeostasis has been identified as a critical factor in male infertility (19), as spermiogenesis is associated with dramatic redox transitions leading to widespread oxidation of thiols. Consistent with this notion, sperm is particularly susceptible to reactive oxygen species (ROS) during critical phases of sperm development (20). TXNRD3 is one of most abundant thiol oxidoreductases during sperm maturation, so its deficiency may be particularly detrimental for the thiol redox transition. By using CM-H2DCFDA to detect ROS, we found no significant differences between KO and WT sperm (Fig. 4*A*). This is not unexpected - while intracellular thiols are among the major targets of ROS (21), thiol redox dysfunction does not necessarily have to involve ROS. To assess potential protein thiol changes in the *Txnrd3* KO sperm, we employed BODIPY-NEM as a fluorescent probe to directly label thiols in sperm samples (22), followed by flow cytometry. We observed a lower percentage of BODIPY-NEM positive sperm in *Txnrd3*^-/-^ than in WT mice (WT 86.57±6.12 vs. *Txnrd3*^-/-^ 64.17±9.12, p=0.024, n=3) (Fig. 4*B*), consistent with the idea that TXNRD3 supports the formation or maintenance of thiols during sperm maturation.

**Fig. 4.**
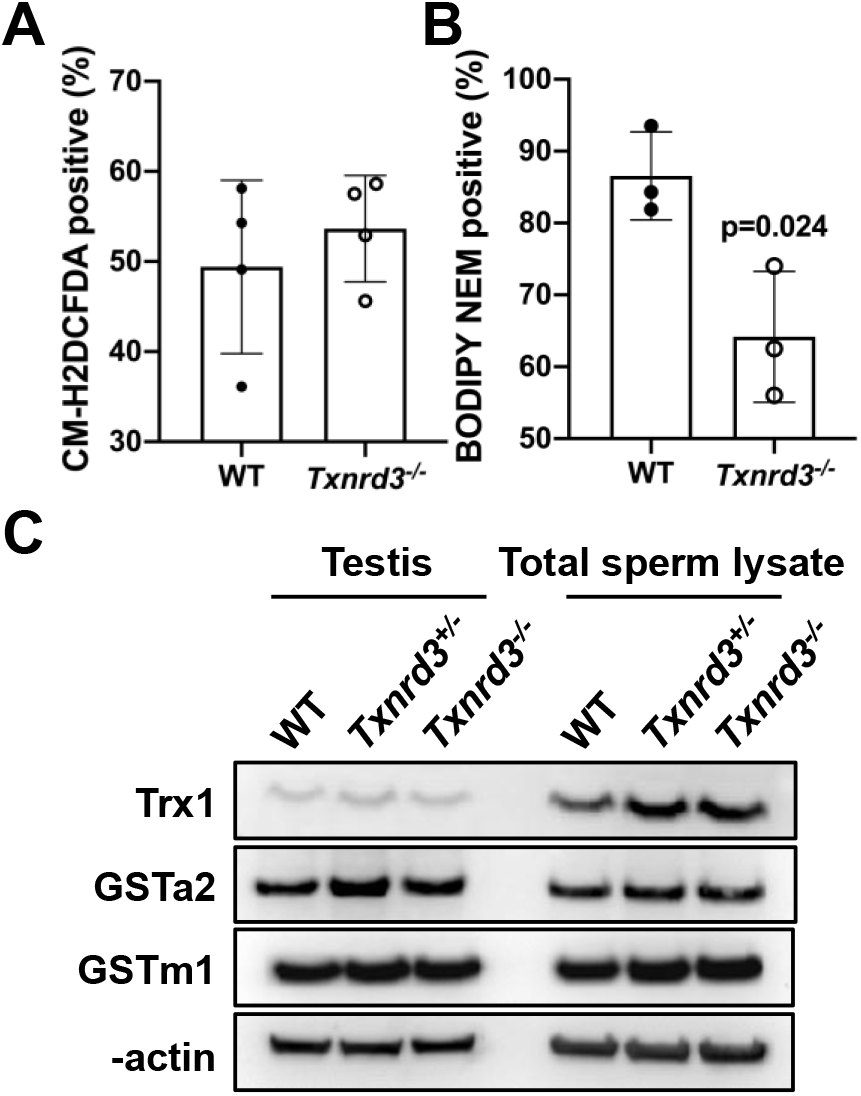
Analyses of redox characteristics of sperm. (A) ROS levels in mouse sperm as revealed by CM-H2DCFDA assays. (B) Protein thiols in mouse sperm as revealed by using BODIPY-NEM. (C) Expression of Trx1, GSTa2, and GSTm1 in mouse testis and sperm by Western blotting.

The most abundant thiol in animal cells is glutathione (GSH), which is present both in the cytosol and various organelles (23). We examined GSH levels and found no differences between WT, *Txnrd3* ^+/-^, and *Txnrd3* ^-/-^ mice (Fig. 5*A*). In addition, we preformed immunoprecipitation to enrich glutathionylated proteins in mouse testis and sperm, and again found no differences among animal groups (Fig. 5, *B* and *C*). We also tested thioredoxin 1 (Trx1), glutathione S-transferase A2 (GSTa2), and glutathione S-transferase Mu 1 (GSTm1) expression in the testis and sperm (Fig. 4*C*). The expression of Trx1 in *Txnrd3*^-/-^ sperm was elevated. This is a conserved thiol oxidoreductases that maintains protein thiols in the reduced state; it is also a substrate for TXRNDs, including TXNRD3 (24). The increased Trx1 expression in *Txnrd3*^-/-^ sperm and testes might be a compensatory mechanism in view of TXNRD3 deficiency.

**Fig. 5.**
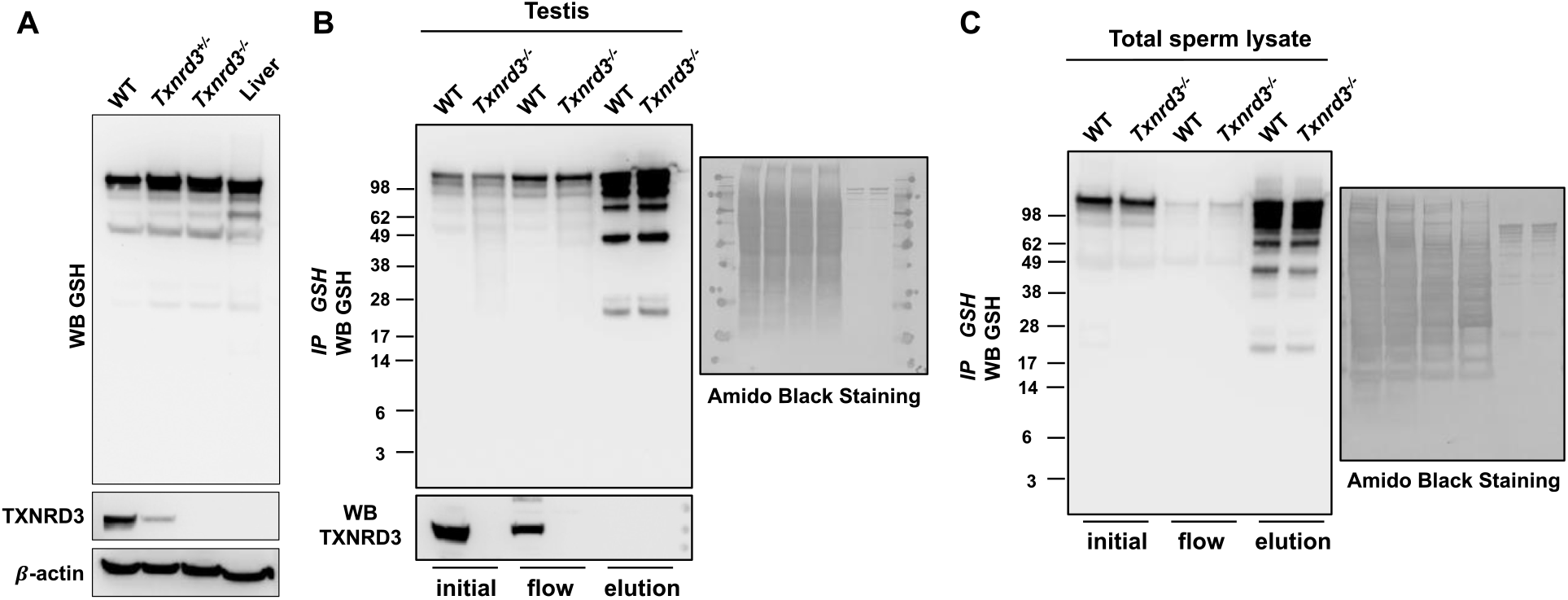
Analyses of glutathionylation in mouse testis and sperm. (A) Glutationylated proteins in the testis of WT and *Txnrd3* KO mice (and liver as control) as analyzed by Western blotting with anti-GSH antibodies. (B) Immunoprecipitation (IP) of glutathionylated proteins in WT and *Txnrd3* KO testes with anti-GSH antibodies. Initial (corresponds to whole cell lysate), flow through (flow) and elution (elution) factions were analyzed by Western blotting using anti-GSH and TXNRD3 antibodies (*left panel*). Protein staining with Amido Black (*right panel*). (C) Immunoprecipitation (IP) of glutathionylated proteins in WT mice and *Txnrd3* KO sperm with anti-GSH antibodies. Initial (corresponding to whole cell lysate), flow through (flow) and elution (elution) factions from a GSH enrichment experiment were analyzed by Western blotting using anti-GSH and TXNRD3 antibodies (*left panel*). Protein staining with Amido Black (*right panel*).

During spermiogenesis, the sperm DNA becomes highly compact. To facilitate this extreme transition, the vast majority of somatic histones are replaced by small basic proteins, and then these proteins undergo thiol oxidation to form intramolecular and intermolecular disulfide bonds (25). These disulfides are crucial for DNA packaging, as its mispackaging in the sperm head results in decreased fertility, higher rates of miscarriage and higher rates of genetic disease in the offspring (26). To test the possibility that TXNRD3 deficency affects disulfide bond formation, we employed diagonal gel eletrophoresis, thereby examining oligomeric proteins containing interchain disulfide bonds. Overall protein patterns of both WT and *Txnrd3*^-/-^ testes and sperm showed similar patterns of protein crosslinking (Fig. S3). Our findings suggest dysfunction in thiol redox homeostasis might occur during spermiogenesis and sperm maturation in *Txnrd3*^-/-^ males.

### Putative TXNRD3 targets

As TXNRD3 exhibits NADPH-dependent thiol reductase activities *in vitro*, and its deficiency reduces the levels of thiols in sperm, a reasonable possibility is that this enzyme supports the reduction of disulfide-containing substrates during sperm maturation. Aiming to detect such endogenous TXNRD3 targets, we developed a procedure for their enrichment and applied it to three different portions (caput, corpus and cauda) of the epididymal luminal contents including sperm of *Txnrd3* KO and WT mice (8 mice per group). This procedure was based on the assumption that target proteins are reduced by TXNRD3 in WT mice but are partially in the oxidized form in *Txnrd3* KO mice (Fig. S4). Thus, free thiols in the sperm lysates of *Txnrd3* KO and WT mice were first blocked with NEM, and the lysates were applied to a column containing the Grx domain of TXNRD3, which was in the pre-reduced state (corresponding to the NADPH-reduced state of TXNRD3). The targets reduced by this domain were then trapped by using a thiol-trapping column and eluted with DTT. The resulting eluates enriched for TXNRD3 targets were treated with IAM to block their free thiols and subjected to proteomics analyses. Spectral counts of detected proteins (Fig. S5) were normalized using the Normalized Spectral Abundance Factor approach (27, 28). Interestingly, we detected more proteins in KO than in WT samples, consistent with the broad range of substrates. To uncover relevant patterns of proteins present in *Txnrd3* KO but not in WT samples, we devised four selection criteria (Fig. S6), and their application to the proteomic dataset led to a set of putative target proteins (Fig. 6). Interestingly, this approach did not reveal a single bona-fide TXNRD3 target. Instead, we detected a broad range of target proteins. Based on functional enrichment analysis (DAVID functional annotation tool v6.8), the set of putative targets (103 proteins) was enriched for proteins annotated as RNA-binding (p-value=1.77E-5; FDR=0.024). In addition, a set of mitochondrial proteins (10 proteins) involved in metabolism were detected only in the cauda sperm from *Txnrd3* KO mice, suggesting that TXNRD3 deficiency might affect ATP production by sperm cells. This finding is consistent with the idea that TXNRD3 reduces a broad set of proteins during spermiogenesis and sperm maturation.

**Fig. 6.**
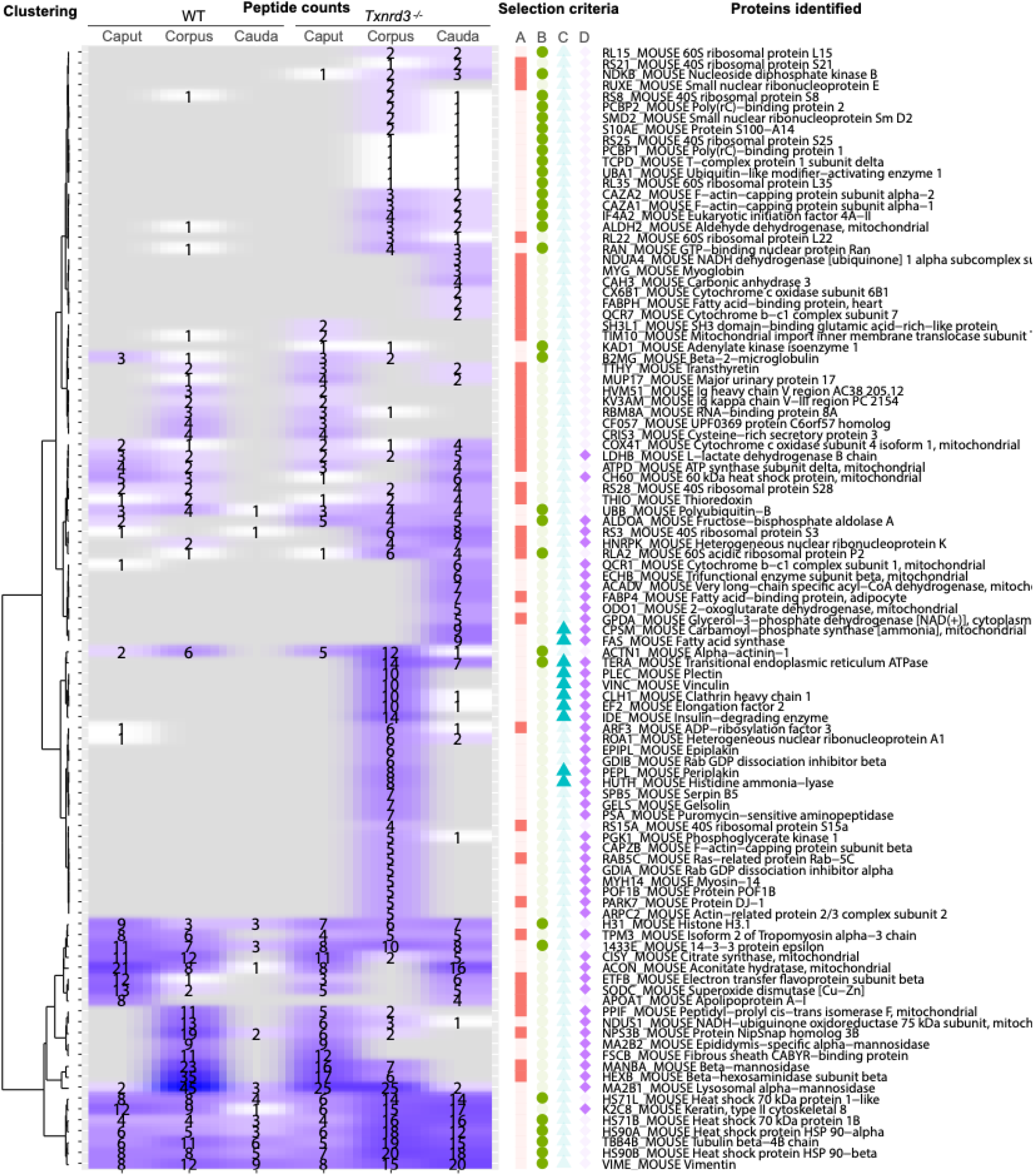
Putative TXNRD3 target proteins. Heatmap shows absolute peptide counts per protein. The left dendrogram represents a clustering structure of candidate proteins (obtained by the R function hclust, ward.D method, applied to the Euclidean distance matrix). On the right of the heatmap, colored shapes show, for each protein, to which candidate set(s) it belongs to (Fig. S4). Finally, the name of candidate proteins is reported on the right; those involved in RNA binding are in bold.

## Discussion

Both previously characterized mammalian TXNRD family members, *i*.*e*., cytosolic TXNRD1 and mitochondrial TXNRD2, are essential proteins. As the third member of this family, TXNRD3 (TGR), is abundantly expressed in the testes across tetrapods, we considered a possibility that this protein instead plays an essential role in spermatogenesis. We developed *Txnrd3* KO mice and, as predicted, found that they were viable and without gross phenotypes. Unexpectedly, however, *Txnrd3* KO males were fertile, *i*.*e*., TXNRD3 was not essential for male reproduction. However, by applying a mating scheme to more carefully examine overall fertility of WT and KO animals, we found that *Txnrd3*^-/-^ males may be subfertile. Additionally, an IVF experiment showed a lower fertilization rate of *Txnrd3*^-/-^ mice compared to WT mice, and the *Txnrd3*^-/-^ sperm exhibited an altered state of its thiols. These data revealed that, while TXNRD3 is not essential for male reproduction, it plays an important role in this process by maintaining redox homeostasis during spermiogenesis. As TXNRD3 is conserved in mammals, presumably its function is also conserved, and the subfertility associated with its deficiency must be severe enough in the wild to preserve the protein in these organisms over evolutionary timescales.

Due to is domain organization, TXNRD3 may be viewed as a component of both Trx and GSH systems. The Trx system provides reducing equivalents to many proteins, but especially to thiol peroxidases, such as PRDXs. It was found that Trx1 can be directly reduced by TXNRD3; this TXNRD3 could support the reduction of thiol peroxidases via this thioredoxin (29, 30). Additionally, Trx1 expression was higher in *Txnrd3*^-/-^ mice sperm (compared to WT mice), suggesting a possible compensatory mechanism. One of PRDXs, PRDX4, was also implicated in male reproduction through redox homeostasis (31), and yet another thiol peroxidase that is a key redox regulator of male reproduction is GPX4 (32). Recently, it was shown that GPX4 expression is increased in the testis and caput epididymis of *Prdx4* knockout mice (33). In addition, our previous study showed that upon prolonged selenium-deficiency, the expression of TXNRD3 and GPX4 is markedly decreased in testes (7). In our current study, we found that GPX4 expression was increased in *Txnrd3*-deficient testis, whereas PRDX4 expression was decreased.

During spermiogenesis, most somatic histones are replaced by small basic nuclear proteins called protamines that condense the spermatid genome into a transcriptionally inactive state (34, 35). Protamines contain multiple cysteine residues, which undergo thiol oxidation to first form intramolecular, and then intermolecular disulfide bonds during epididymal maturation. These disulfide bonds stabilize the sperm DNA and are thought to be crucial for condensing the mammalian sperm nucleus into its fully mature state (26). While sperm maturation is associated with thiol oxidation, TXNRD3 is a reductase. It was previously suggested that TXNRD3 may catalyze isomerization of disulfide bonds (7) as well as the reduction of incorrectly formed disulfide bonds. Both of these functions are consistent with the data generated with *Txnrd3* KO mice, and especially the latter one. Indeed, our BODIPY-NEM findings show a more oxidized state of *Txnrd3*^-/-^ sperm than in WT mice.

If the function of TXNRD3 is to reduce disulfides during spermiogenesis when these disulfides are formed, what could be the TXNRD3 targets? To identify these proteins, we developed an enrichment protocol, whereby these proteins, which were expected to be more oxidized in the *Txnrd3* KO sperm, were reduced on the column containing a recombinant Grx domain of TXNRD3, and the resulting thiols were trapped by the thiol-trapping column. The resulting enriched proteins were identified by mass-spectrometry. This procedure yielded many putative targets of TXNRD3 as opposed to a single protein. In particular, TXNRD3 targets were enriched for RNA-binding proteins (RBPs) and mitochondrial proteins. RBPs are abundantly expressed throughout spermatogenesis, and are reliant on mechanisms of posttranscriptional regulation of gene expression (36). Knockout of genes encoding specific RBPs or testis-expressed RBP disruption often cause an irregular spermatogenesis phenotype, various stages of spermatogenic arrest, and male infertility, suggesting that each of the steps of post-transcriptional regulation during spermatogenesis needs to be finely tuned by specific proteins to ensure the production of fertile male gametes (36, 37). For example, RNA-binding protein 8A (Rbm8a) was detected in our protein enrichment analysis, and was highly expressed in spermatogonia and spermatocytes, while had low levels in late spermatids. Rbm8a, an exon junction complex protein, plays an important role in regulation of translation, splicing, RNA transport and localization. Genetic ablation of Rbm8a can cause early embryonic lethality (38, 39). Our data support the idea that, in *Txnrd3*^-/-^ mice, incorrectly formed disulfide bonds in RBPs may cause protein dysfunction, leading to subfertility.

Mitochondria participate in diverse processes in support of spermatogenesis and fertilization (40). Sperm cell motility is dependent on ATP production (41). 2-oxoglutarate dehydrogenase (OGDH, ODO1), citrate synthase (CS, CISY), ATP synthase subunit delta (ATPD), and cytochrome c oxidase subunit 4 isoform 1 (COX4-1), which are potential TXNRD3 targets, are involved in ATP production (42, 43). A decrease in OGDH has been linked with reduced sperm motility (44). Cytochrome b-c1 complex subunit 1 (UQCRC1, QCR1) is associated with oxidative stress (45), and its UQCRC1 expression correlates with litter size (46, 47). Aldehyde dehydrogenase (ALDH2) is involved in intracellular sperm antioxidant defense system and plays a pivotal role in the maintenance of sperm motility by removing 4HNE (a product of lipid peroxidation) (48), an acetaldehyde that could damage the sperm DNA (49). The 60 kDa heat shock protein (HSPD1, CH60) is involved in sperm capacitation through tyrosine phosphorylation (50). The detection of these potential TXNRD3 target proteins suggested a possibility that TXNRD3 supports a mitochondrial function during sperm capacitation. In the accompanying manuscript, we provide further evidence for this possibility as well for the key role of TXNRD3 in redox regulation during epididymal maturation, which underlies capacitation-associated mitochondrial activation and sperm motility (16).

Overall, we show that *Txnrd3* KO is associated with reduced male fertility both *in vitro* and *in vivo* due to altered thiol redox homeostasis. This phenotype is associated with the inability to reduce incorrect disulfides formed during the widespread oxidation of thiols in process of sperm maturation.

## Acknowledgments

This work was funded by NIH R01GM065204 to V.N.G, and the start-up funds from Yale School of Medicine and NIH R01HD096745 to J.-J.C.

