## Supplementary Information for "TXNRD3 supports male fertility via the redox control of spermatogenesis"

### Supplementary Figure S1

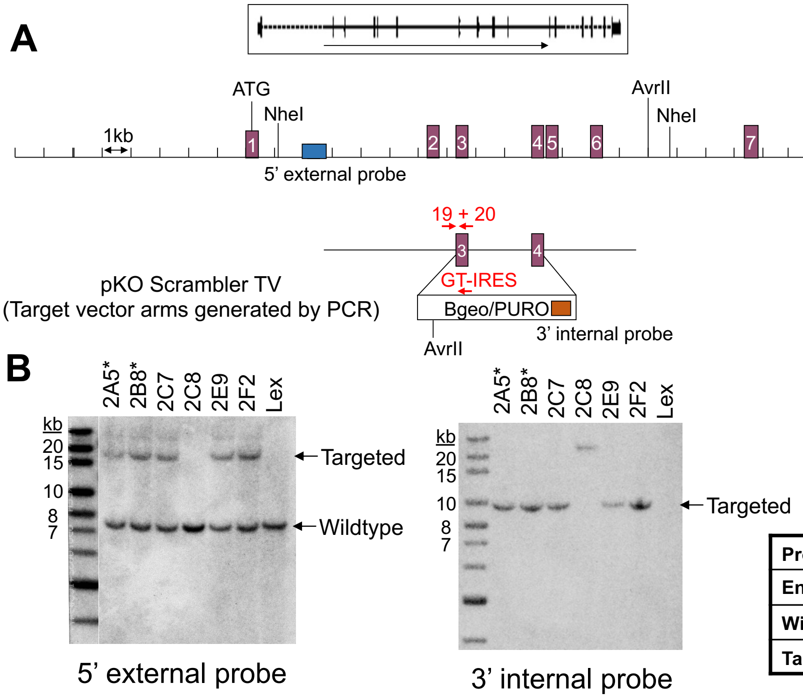


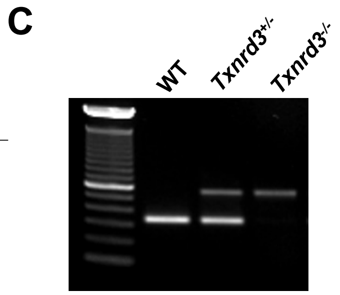


| Strategy | WT-specific | Mutant-specific |
|---|---|---|
| Primers | 19+20 | 19+GT-IRES |
| Wildtype | 322 bps | ----------- |
| Targeted | -------- | 441 bps |

| Probe | 5’ external | 3’ internal |
|---|---|---|
| Enzyme | NheI | AvrII |
| Wildtype | 7.6 kb | ------- |
| Targeted | 17.4 kb | 9.8 kb |

**Fig. S1. Generation of *Txnrd3* knockout mice**. (A) Targeting strategy by homologous recombination in ES cells. Targeting vector is pKO Scrambler TV. (B) Southern blot confirmation of the homologous recombinant ES cell clones. Hybridization with the 5’ external probe showing a 7.6 kb band specific to the wild type allele, and a 17.4 kb band specific to the targeted allele. Hybridization with the 3’ internal probe showing a 9.8 kb band specific to the targeted allele. (C) Identification of *Txnrd3*-deficient mice by PCR. Wild type (WT) mice were distinguished by PCR products from primer set 19+20, null mice (-/-) were positive for 19+GT-IRES, and heterozygous mice (+/-) were positive for both.

**Supplementary Figure S2**

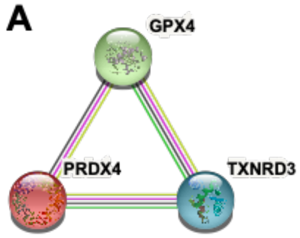


| Node1 | Node2 | Neighborhood_on_chromosome | coexpression | Experimentally_determined_interaction | Automated_textmining | Combined_score |
|---|---|---|---|---|---|---|
| GPX4 | TXNRD3 | 0.172 | 0.151 | 0.135 | 0.59 | 0.717 |
| GPX4 | PRDX4 | 0 | 0.062 | 0.197 | 0.629 | 0.696 |
| PRDX4 | TXNRD3 | 0.161 | 0.117 | 0.122 | 0.383 | 0.544 |

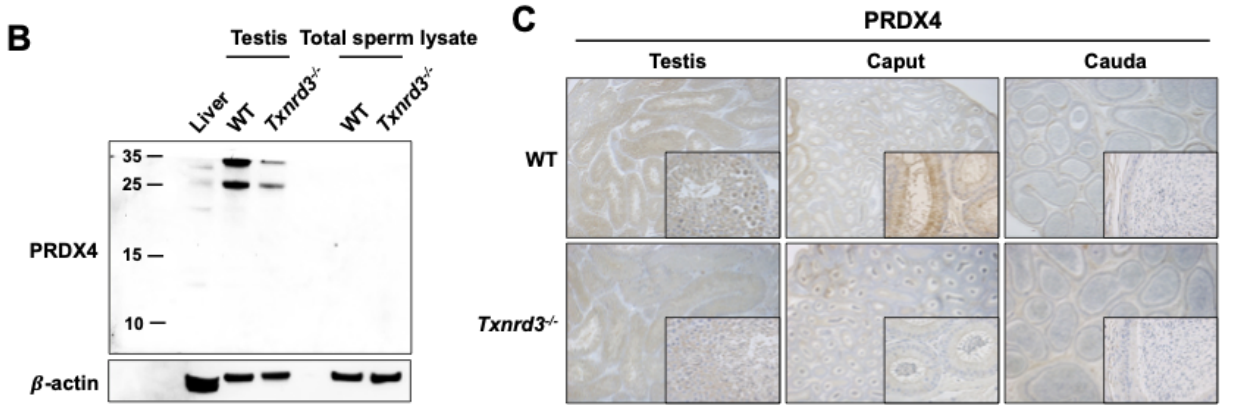


**Fig. S2. Functional associations involving TXNRD3, PRDX4 and GPX4 in mouse testis and sperm**. (A) Association types in STRING, shown separately for each data source and confidence range (low confidence: scores <0.4; medium: 0.4 to 0.7; high: >0.7) (51). (B) PRDX4 expression in mouse testis and sperm by Western blotting. (C) PRDX4 in mouse testis and sperm (caput and cauda) as analyzed by immunohistochemistry.

#### Supplementary Figure S3

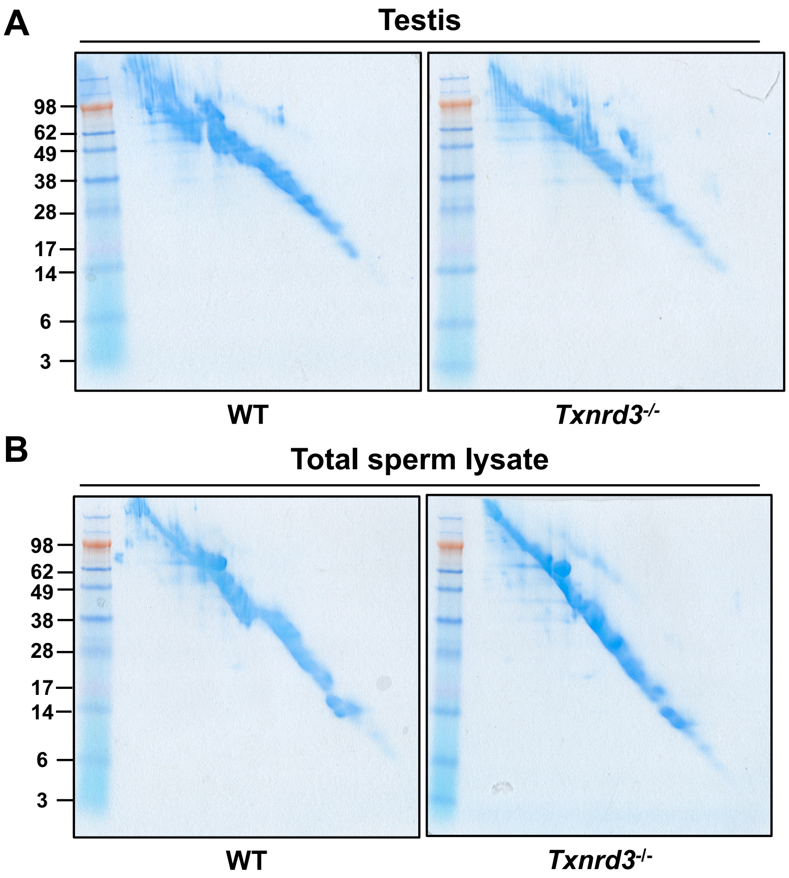


**Fig. S3. Detection of mixed disulfide complexes in testis and sperm.** Diagonal two-dimensional analyses were performed on mouse testis lysate (A) and total sperm lysate (B), where the first non-reducing electrophoresis was followed by reducing electrophoresis in the second dimension.

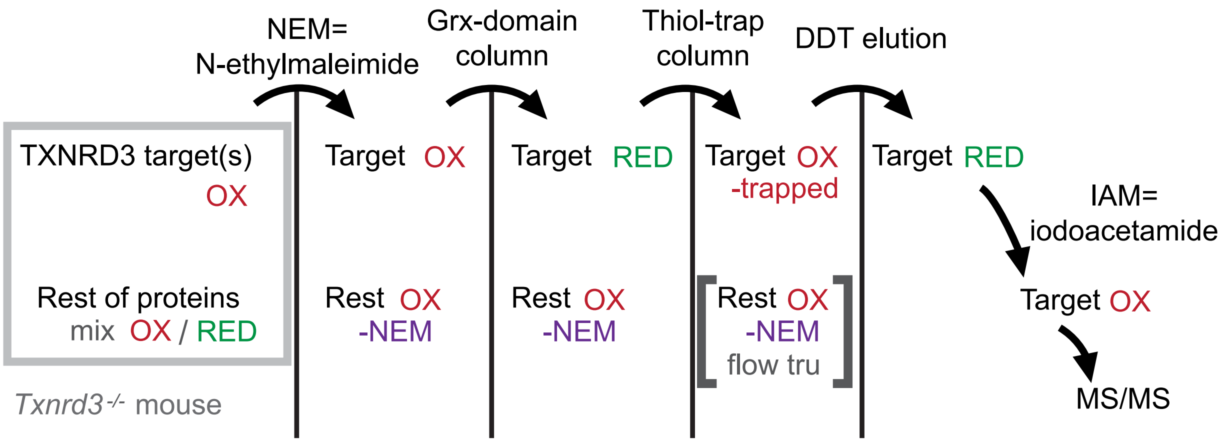


**Fig. S4 Experimental design of the experiment to identify putative TXNRD3 target proteins in the sperm of *Txnrd3* KO mice.** Shown are target proteins and other proteins at each step of the enrichment procedure. Proteins in WT mice are not shown but were used as a control in the experiment. Lysates of sperm (separately for cauda, corpus, caput fractions) were treated to NEM to block free thiols and subjected to a column containing a pre-reduced recombinant Grx domain of TXNRD3. The proteins reduced by this domain were then trapped by a thiol-trapping column, eluted with DTT, protected by IAM and analyzed by MS/MS.

#### Supplementary Figure S5

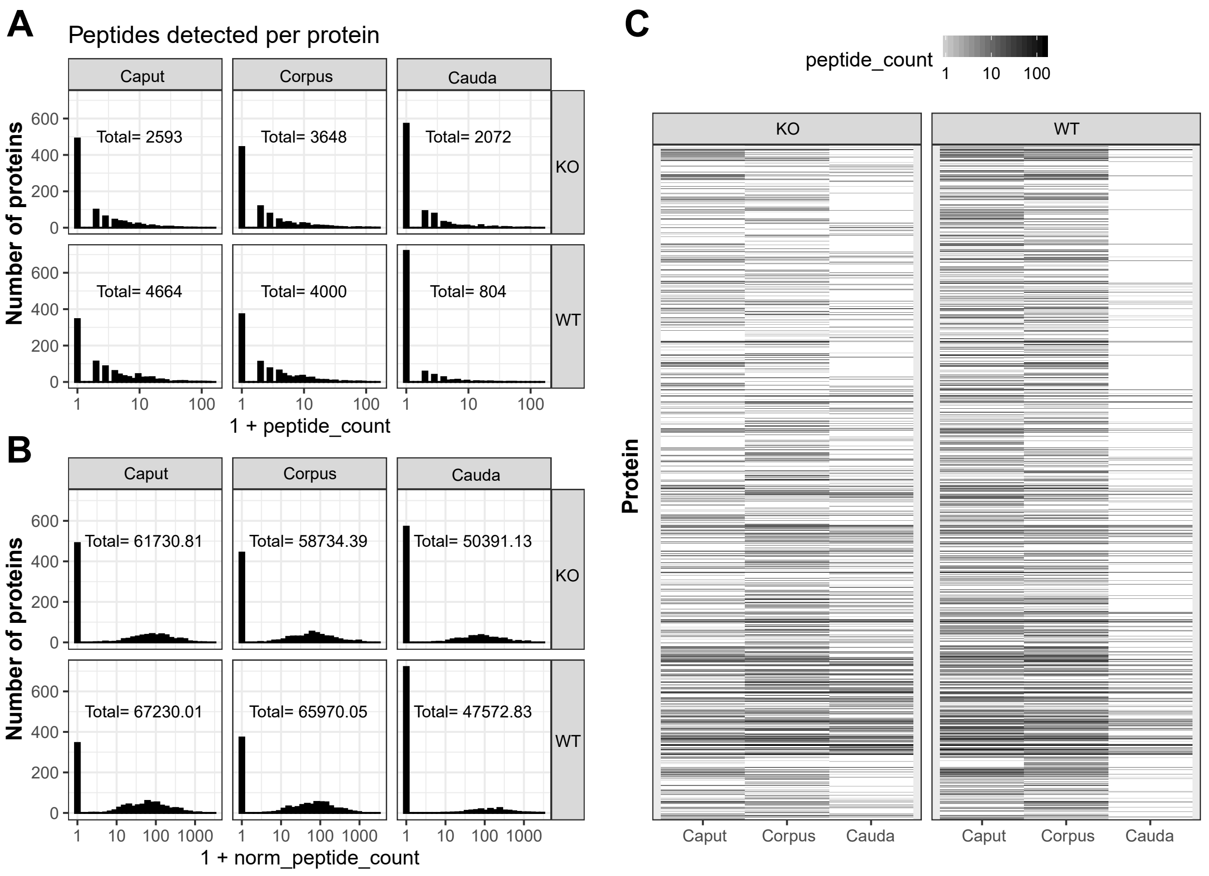


**Fig. S5. Overview of LC-MS/MS data**. (A) Distribution of the number of peptides detected per protein (absolute counts). (B) Distribution of normalized peptide counts. (C) Heatmap of absolute peptide counts per protein.

#### Supplementary Figure S6

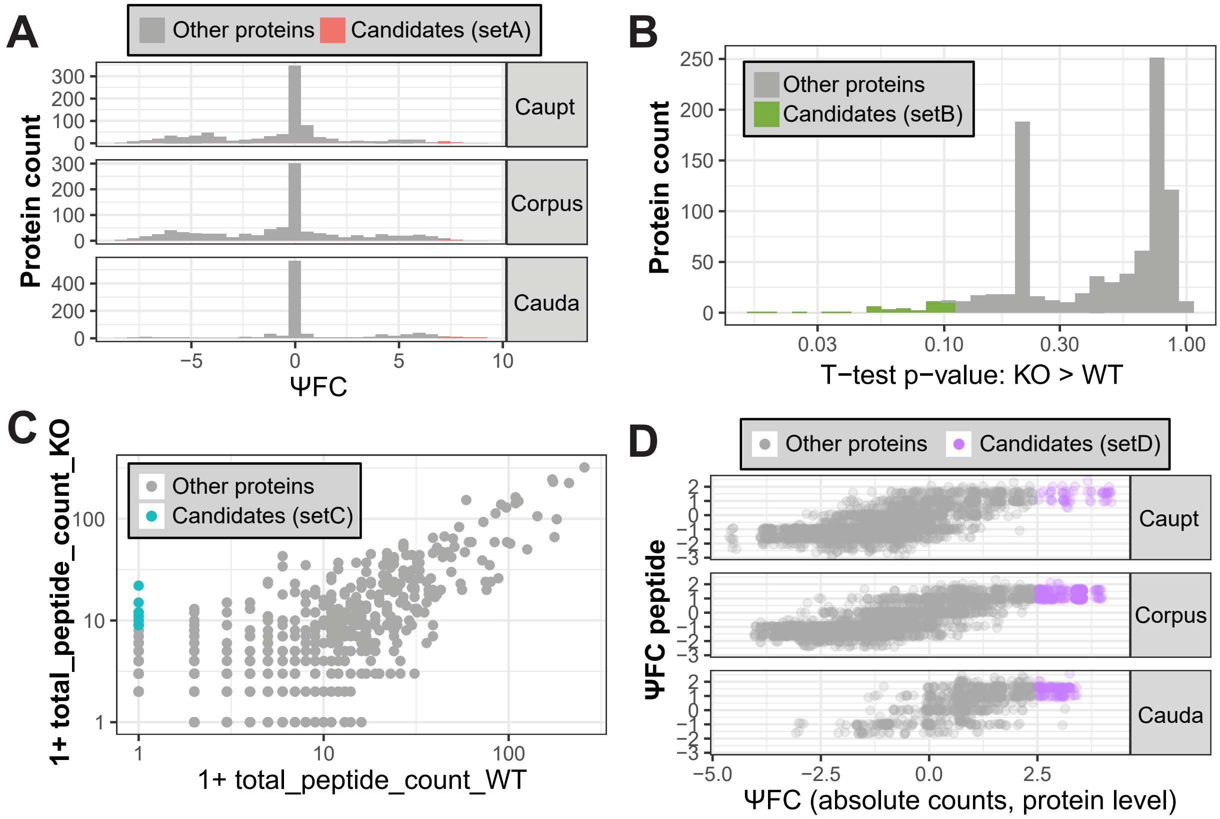


**Fig. S6. Overview of the four selection criteria and resulting sets of candidate proteins**. (A) We defined a metrics, analogous to log fold-change with pseudocounts, called ΨFC:

$$\Psi FC = \log 2\left(\frac{(KO + 1)}{(WT + 1)}\right)$$

wherein KO and WT are the normalized counts in KO and WT samples, respectively. We computed ΨFC for each protein in each testis fraction (caput, corpus, and cauda). We considered all proteins with ΨFC>=7 in any fraction as candidates (setA). (B) We treated caput, corpus, and cauda as replicates, and performed a one-tailed paired T-test. We considered all proteins with nominal p-value <10% as candidates (setB). (C) We considered as candidates (setC) all those proteins with no absolute peptide counts in any of the WT samples, and at least 8 counts in total in KO samples. (D) We combined quantifications at protein and single peptide levels, and we considered as candidates (setD) those proteins with ΨFC > 2.5 and with any peptide whose $\Psi FC_{peptide} > 0$ in any testis fraction (where $\Psi FC_{peptide}$ is analogous to ΨFC as defined above, but considering peptide counts instead).

#### Supplementary Table S1

| | *Txnrd3*$^{+/-}$ x *Txnrd3*$^{+/-}$ | | | | |
|---|---|---|---|---|---|
| | WT | *Txnrd3*$^{+/-}$ | *Txnrd3*$^{-/-}$ | Total[1] | Ratio[1] |
| Female | 32 | 49 | 18 | 99 | 1 |
| Male | 30 | 56 | 17 | 103 | 1.04 |
| Total[2] | 62 | 105 | 35 | 202 | |
| Ratio[2] | 1 | 1.7 | 0.56 | | |

**Table S1. Genotypes of offspring from *Txnrd3*$^{+/-}$ x *Txnrd3*$^{+/-}$ crosses.** This table shows the number of pups in each group. Total[1]: total number of female mice and male mice; Total[2]: total number of WT mice, *Txnrd3*$^{+/-}$ mice and *Txnrd3*$^{-/-}$ mice; Ratio[1]: ratio of female mice and male mice; Ratio[2]: ratio of WT, *Txnrd3*$^{+/-}$ and *Txnrd3*$^{-/-}$ mice.

#### Movie S1

**Movie S1. Movement of free-swimming spermatozoa from WT, *Txnrd3*$^{+/-}$, and *Txnrd3*$^{-/-}$ mice.** Uncapacitated spermatozoa were allowed to disperse for 15 min pre-incubation in a 37 °C containing M2 media; free swimming sperm cells recorded within the next 5 min; 2s movies.
